# Between- and within-host mutation of dengue virus

**DOI:** 10.1101/2025.10.12.680595

**Authors:** Sachith Maduranga, Braulio Mark Valencia, Chathurani Sigera, Praveen Weeratunga, Deepika Fernando, Senaka Rajapakse, Melissa Rapadas, Ira W. Deveson, Andrew R Lloyd, Rowena Bull, Chaturaka Rodrigo

## Abstract

RNA viruses exhibit high mutation rates due to error-prone polymerases, leading to a diverse pool of viral haplotypes (also referred to as quasi-species) within infected hosts. While haplotypes have been well studied in chronic infections like HIV and HCV, diversity remains under-explored in acute infections like dengue (DENV), which are constrained by a short viremic phase. This study aimed to characterise the mutation hotspots in DENV genomes at both consensus and haplotype levels.

Near full length DENV genomes were sequenced using Oxford Nanopore Technology (ONT) from the plasma of Sri Lankan patients with dengue fever recruited between 2017 -2020. Consensus sequences were mapped with Minimap-2, and haplotypes were reconstructed with Nano-Q, a tool designed for estimation of RNA virus haplotypes and their relative abundance. The genomic variability of DENV genomes was assessed by calculating Shannon Entropy (SE). Codons undergoing diversifying selection were identified with three phylogenetics-based algorithms (FEL, MEME, FUBAR) implemented within the Datamonkey suite.

From 150 samples tested, both consensus and haplotype sequences were characterised in 90 samples (DENV1: 8, DENV2: 51, DENV3: 31). The genomic variability of consensus sequences measured by SE was higher in DENV2 compared to DENV3, and the reverse was true for haplotypes. At the consensus level, the NS2A gene had the greatest number of mutable sites when adjusted for gene length across all serotypes, while at the haplotype level the NS1 gene had the same. Overall, the haplotypes sequences revealed more sites with high mutability and codons under diversifying selection than those visible at consensus level. This provides proof-in-principle that in acute RNA viruses also have high mutability in haplotypes, which may be inapparent with a consensus-level analysis.

## 1. Introduction

RNA viruses exist as a population of genomically different variants or haplotypes (within-host variants, quasi-species) in an infected host due to the lack of fidelity in RNA-to-RNA replication (Domingo & Perales, 2019; Rodrigo & Luciani, 2019). The haplotypes and their relative abundance is continuously changing in response to viral fitness and evolutionary pressures exerted by the host immune system and antiviral treatment (if available) (Rowena A. Bull et al., 2011). On occasion, more fit haplotypes sweep the quasi-species population when encountered with formidable selection pressures known as genetic bottlenecks (e.g., after initiation of antiviral treatment, or upon transmission of infection to a new host) (Rowena A. Bull et al., 2011; Zhang et al., 2021). Genetic bottlenecks are well characterised in chronic infections like Hepatitis C virus (HCV) and Human Immunodeficiency virus (HIV) infections (Forns, Purcell, & Bukh, 1999). In such instances, it is not uncommon for a low-abundant haplotype (minor variant) to become dominant in the quasi-species population if it carries fitness mutations to allow better survival at the bottleneck. Thus, characterising haplotypes, even those with low abundance is important to predict disease outcomes and treatment response in RNA virus infections.

In contrast to chronic RNA virus infections, acute infections (e.g., SARS-CoV-2, influenza, dengue) are short-lived and typically cleared by the human immune system in a matter of days. The shorter period of viraemia limits continuous within-host virus evolution in these infections. Nevertheless, these viruses causing acute infections also mutate, survive and transmit at community level very efficiently as demonstrated by periodic epidemics or even pandemics (Hu, Guo, Zhou, & Shi, 2021). Within-host viral evolution is the stepping stone for such efficient between-host evolution, but characterising the former in acute RNA virus infections has not been at the same forefront as for chronic RNA viruses. Characterising within-host mutation hotspots in acute infection causing RNA viruses can inform specific regions to be targeted in genomic surveillance of outbreaks, in vaccine research and for surveillance of emerging resistance to antivirals.

Dengue fever is an acute RNA virus infection with an estimated incidence of 390 million infections globally, including asymptomatic transmissions (Bhatt et al., 2013). With half the global population at risk of infection, it is an emerging threat to global health. While most symptomatic patients recover fully from a flu-like acute illness within 2-weeks, a minority progress to severe dengue characterised by bleeding, severe plasma leakage (extravasation of plasma into interstitial space) with shock, and organ dysfunction (Rajapakse, Rodrigo, & Rajapakse, 2012; World Health Organization, 2009). Dengue virus (DENV) has a positive sense RNA genome which is approximately 10,600 nucleotide-long and its coding region is translated as a 3386 amino acid-long single polyprotein (T. N. Adikari et al., 2020) split into 3 structural [Capsid (C). envelope (E), membrane (M)] and 7 non-structural (NS1, NS2A, NS2B, NS3. NS4A, NS4B, NS5) proteins. DENV has four serotypes (DENV1-4) further subdivided by phylogenetics into 19 genotypes based on distinct differences in genetic code (Yung et al., 2015).

Haplotype characterisation of RNA viruses had been constrained by the technical challenges in isolating uninterrupted viral genomes and sequencing them in-whole, without fragmentation. With the introduction of whole virus genome amplification assays and third generation long-read sequencing platforms (e.g., Oxford Nanopore Technology – ONT) these challenges are gradually being resolved (Rodrigo & Luciani, 2019). Previously we published near full-length virus genome amplification assays for both HCV and dengue viruses, and a bioinformatics tool (Nano-Q) to estimate haplotypes and their relative abundance using ONT sequencing data (T. N. Adikari et al., 2020; R. A. Bull et al., 2016; Riaz et al., 2021; Riaz, Leung, Bull, Lloyd, & Rodrigo, 2022). The present work aimed to characterise the mutability of the dengue virus genome at both between-host and within-host levels using these tools. In this manuscript the term haplotype is synonymous with “within-host variants” or “quasi-species” and will be used throughout as the preferred term for consistency.

## 2. Methods

### 2.1 Clinical samples and viral sequence generation

The clinical samples for this study were acquired from the Colombo Dengue Study (CDS) which is a prospective cohort study (with a nested case-control design) that recruited adult patients with clinically suspected dengue fever presenting within the first 96 hours of fever, from the National Hospital of Sri Lanka in Colombo, Sri Lanka. The purpose, study design, inclusion and exclusion criteria of CDS were published previously (C. Sigera et al., 2021; P. C. Sigera et al., 2019). In brief, the primary purpose of CDS was to identify early risk predictors of dengue associated plasma leakage and ways of improving cost-effective patient management in resource limited settings. Diagnosis of dengue in all CDS recruits was confirmed by a bedside NS1 antigen test and a RT PCR test(Santiago et al., 2013). The patients were admitted and prospectively observed in the hospital until discharge to record demographic, clinical (signs and symptoms on admission and their daily evolution) and the results of laboratory investigation done for routine management. The primary outcome observed in CDS patients was the occurrence of plasma leakage (PL), a preceding event for severe dengue.

The experiments described in this paper used CDS plasma samples collected from all confirmed dengue patients between 2017 October and February 2020. Near-full-length DENV genomes were generated as previously described using a nested PCR assay (Thiruni N. Adikari et al., 2020). The amplicons were visualised on a 0.8% agarose gel, cleaned and size selected using magnetic beads (Ampure®, Beckman Coulter, CA, USA), and quantified with the Quant-iT Picogreen® assay (ThermoFisher, USA, catalogue number: P7589) prior to being sequenced with Oxford Nanopore Technology by the service provider, Garvan Genomics Platform, Sydney, Australia. The sequencing was done on a PromethION flow cell (FLO-PRO114M) with native barcoding (SQK-NBD114-96), multiplexing between 24 – 48 samples in a single run. The base called data were filtered for high quality reads (Mean Q score =9) and demultiplexed with Dorado (7.3.11+0112dde09).

### 2.2 Generation of consensus and haplotype sequences

A detailed phylogenetic analysis from all dengue positive samples of CDS patients recruited between 2017 – 2020 was previously published (Maduranga et al., 2023). However, this analysis was restricted to consensus sequences. As a greater depth of sequencing was needed for haplotype estimation (minimum of 1000, and ideally greater than 10,000 full-length reads per sample), all samples which had a visible band on gel electrophoresis in the 10-11kb range after PCR amplification were re-amplified and re-sequenced as described above to provide a greater sequencing depth per sample. The consensus sequences were generated by aligning reads to a serotype specific reference using Minimap-2 aligner (Version: 2.24) implemented within the Geneious Prime (2024) software suite. The autologous reference obtained from this step was used to realign the raw reads in a second iteration (Figure 1). The consensus generated from this step and the raw read file in .fastq format were used to identify within-host viral variants (haplotypes) using the Nano-Q software tool as published previously (Thiruni N. Adikari et al., 2020; Riaz et al., 2021). This tool outputs the haplotype sequences and their relative abundance as a percentage. Nano-Q does so by using a hierarchical clustering algorithm to generate haplotype sequences at a user-defined length, using the portion of raw reads that matches or exceeds this length. As the length of the reconstructed haplotype increases the number of haplotypes detected decreases, because fewer raw reads meet the length cut-off threshold. Therefore, haplotypes were first reconstructed for 10% of the sample at pre-defined lengths of 10,000, 9000, 8000, 7000 and 6000 and the last was identified as best trade-off point to retain 2 or more haplotypes from the greatest number of samples. Thus, the length of reconstructed haplotypes was fixed at 6000 nucleotides (coding region only, 5’to 3’ direction) with additional settings in the algorithm to create clusters of raw reads that had a Hamming distance threshold equal to or less than 100, and then to generate a consensus sequence per cluster (a haplotype) only if each such cluster had a minimum of 20 reads. The cluster size was taken as a proxy measure of haplotype abundance. For a full description of Nano-Q parameters (T. N. Adikari et al., 2020; Riaz et al., 2021) . Given the poor coverage of reads in the extremes of the genome, first 29 codons of the coding “DNA” sequence in the 5’ end had to be discarded when reconstructing haplotypes.

**Figure 1.**
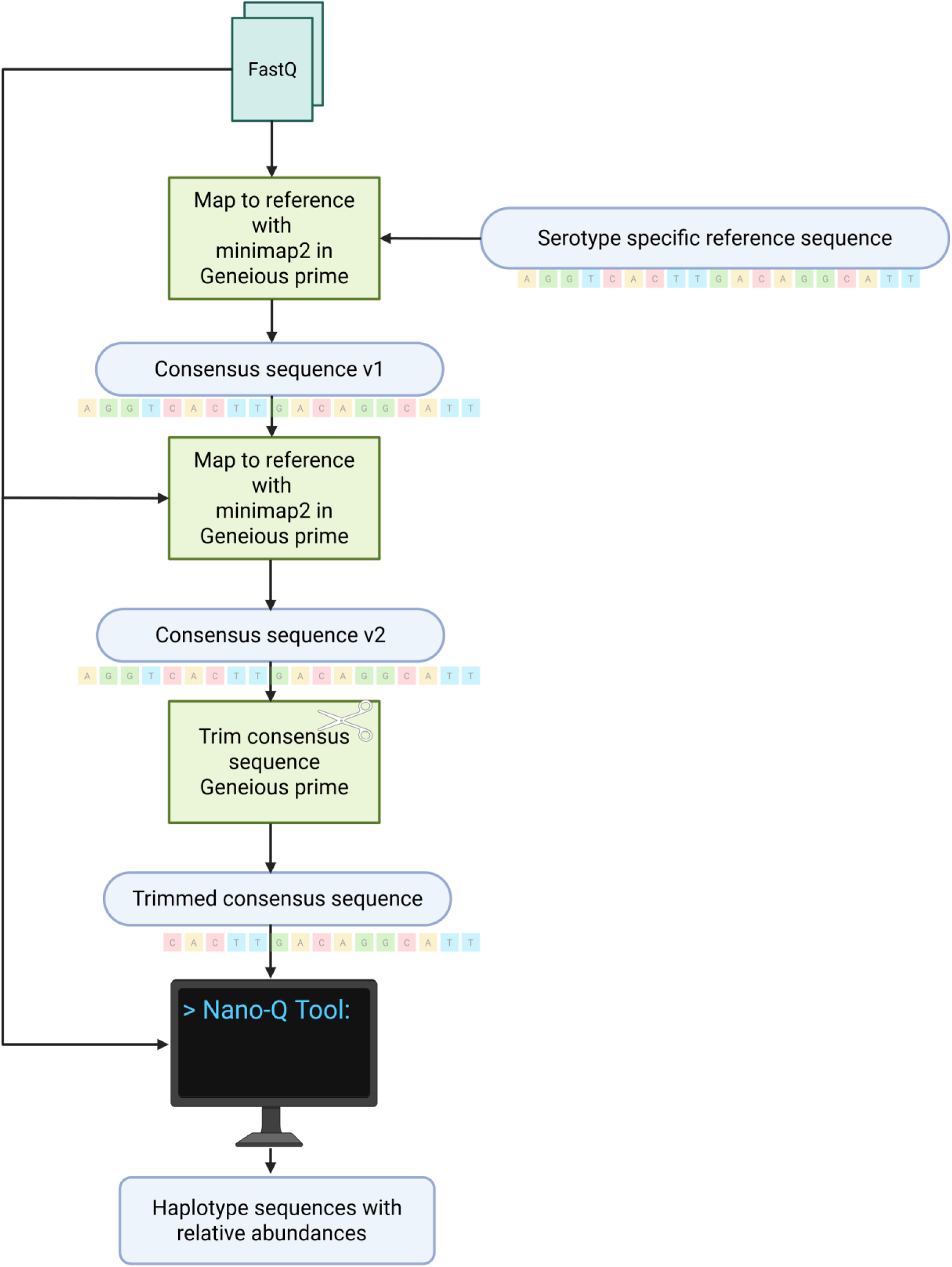
Diagrammatic summary of the generation of the haplotype sequences and relative abundances with the Nano-Q tool. The fastq files were mapped to a serotype specific reference to generate a consensus sequence that was used as the reference for the next round of mapping to reference. The sequences were then trimmed and used as the input for the Nano-Q program with the fastq file. Implementation of Minimap2 (version: 2.24) and trimming of sequences was carried out with Generous Prime (2024). Image created in https://BioRender.com

**Figure 2.**
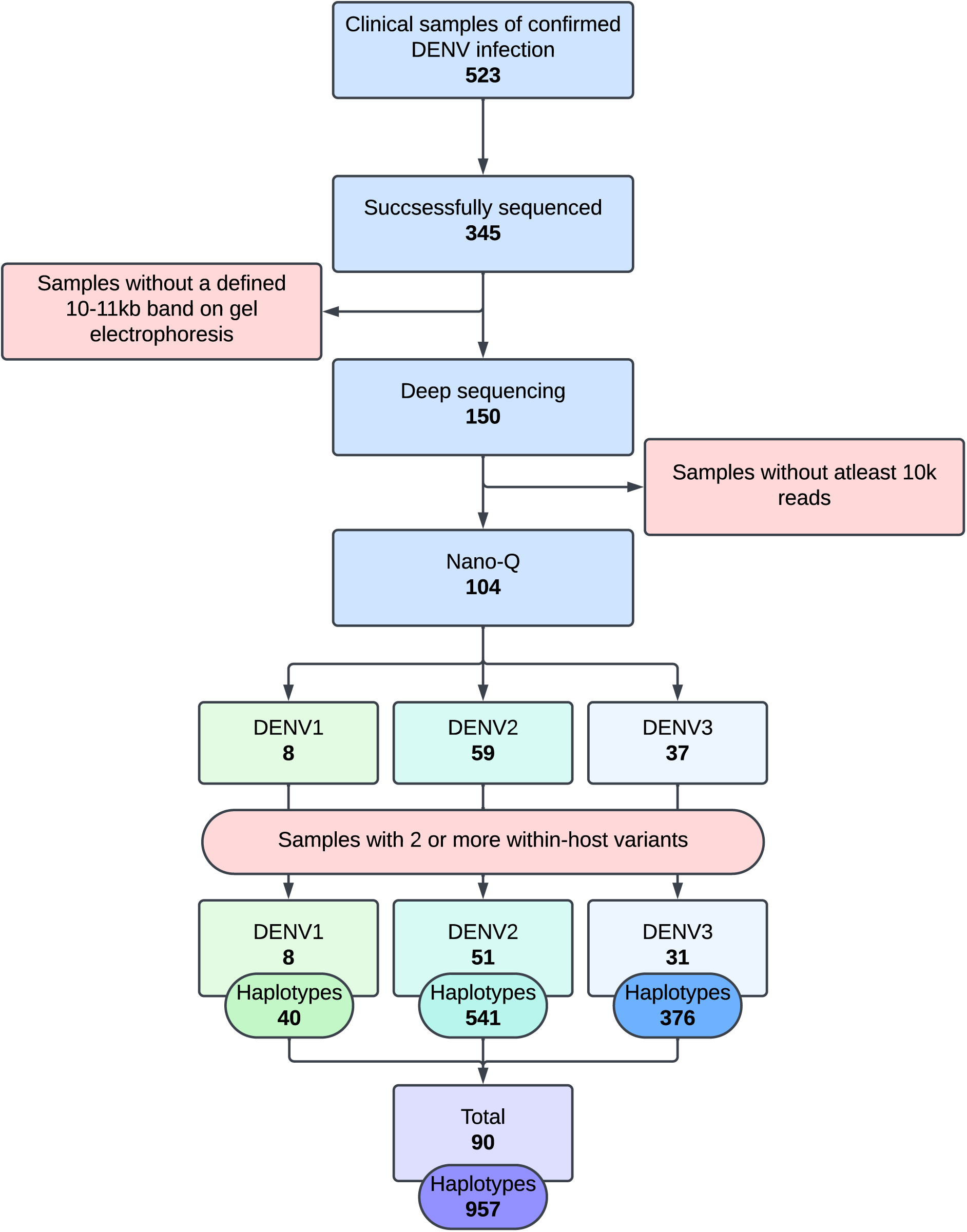
Summary diagram of sample numbers that were analysed at each step. Of the 345 samples that were previously sequenced successfully 150 had adequate post amplification concentration and a defined 10-11kb band in gel electrophoresis. Samples with at least 10k reads were selected for Nano-Q. A total of 90 samples were included in the final analysis with 8 DENV1, 51 DENV2 and 31 DENV3 samples that corresponds to 957 total haplotypes with 40 DENV1, 541 DENV2, and 376 DENV3 haplotypes sequences.

### 2.3 Metrics for mutability of the dengue virus

This analysis was conducted at both consensus (between-host) and haplotype (within-host) levels for each serotype separately. All consensus sequences (approximately 10k nucleotides in length) and corresponding viral haplotypes (6k nucleotides in length) were combined into serotype-specific alignments using the MUltiple Sequence Comparison by the Log-Expectation algorithm (MUSCLE, v3.7), which were then used to calculate the following metrics.

#### 2.3.1. Shannon Entropy (SE)

The variability of non-synonymous mutations in the between-host (consensus) and within-host (haplotypes) sequences were measured with Shannon entropy, a measure of uncertainty (Shannon, 1948). The sequences were translated into amino acids in the correct reading frame and the Shannon entropy was calculated for each codon according to the following formula (equation 1).

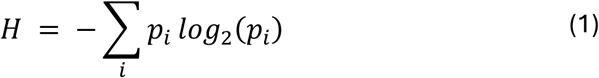

Where, 𝛨 is the Shannon entropy and 𝑝_𝑖_ is the probability of amino acid 𝑖 at a given locus and is calculated as given below (equation 2).

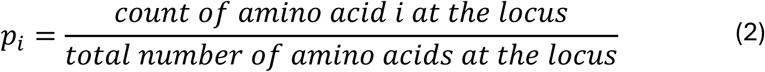

Since 𝑙𝑜𝑔_2_ was used the resulting values are given the unit of “bits”.

The Miller-Madow Bias Correction (de Matos Simoes & Emmert-Streib, 2011; Miller, 1955) was then used to correct for the under-sampling bias associated with calculating Shannon entropy in empirical data (equation 3).

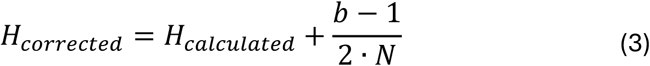

Where, 𝑏 is the number of bins (observed states) and 𝑁 is the sample size.

The standard deviation (SD) of Shannon entropy per serotype specific alignment was calculated and the mean plus twice the SD (Mean + 2SD) was used as a cut-off to identify codons with a significantly higher Shannon entropy.

#### 2.3.2. Codons under diversifying selection

To identify codons undergoing diversifying selection, three algorithms hosted within the Datamonkey online software suite were used in parallel. [https://www.datamonkey.org/ (Delport, Poon, Frost, & Kosakovsky Pond, 2010; Sergei L Kosakovsky Pond et al., 2019; Pond & Frost, 2005; Pond, Frost, & Muse, 2004; Weaver et al., 2018)]. Of these, Fixed Effects Likelihood model (FEL)(Sergei L. Kosakovsky Pond & Frost, 2005) and Fast Unconstrained Bayesian AppRoximation for Inferring Selection (FUBAR)(Murrell et al., 2013) model were used to identify codons with pervasive diversifying selection (p<0.05 and posterior probability > 0.9 respectively), and the mixed effects model of evolution (MEME)(Murrell et al., 2012) was used to identify codons with episodic diversifying selection (p<0.05). These analyses were performed for both within (variant) and between host (consensus) alignments, stratified by serotype. In this paper diversifying hotspots were defined as codons with a SE > mean+2SD and found to have statistically significant diversifying selection by at least one of the above-mentioned algorithms.

For external validation of the findings per serotype, the same analyses were repeated using near-full-length publicly available consensus dengue sequences (DENV1, DENV2, DENV3) from other outbreaks outside of Sri Lanka. For this purpose, DENV sequences were downloaded from the BV-BRC (Bacteria and virus bioinformatic resource centre) database (last date of search: 21 March 2025) guided by the meta-data on the serotype, country and year of isolation and the information available on the linked publications. The same serotype sequences from the same country, in adjacent years linked to a single publication or a project was considered as sampling from a single outbreak. This validation was only done for between-host (consensus) sequences only as equivalent datasets for haplotypes were not available.

## 3. Results

CDS recruited 877 patients between October 2017 and February 2020 of whom 523 were confirmed to have dengue infection by either NS1 antigen testing and/or RT-qPCR. Of these, full genome consensus sequences were successfully amplified from 345 patients, as previously published (Maduranga et al., 2023). Using the gel electrophoresis results from the previous amplification rounds as a guide, a subset of 150 samples with adequate viral RNA were selected for reamplification and resequencing for this project. Following resequencing, consensus sequences were available for all samples while 90 samples (DENV1: 8, DENV2: 51, DENV3: 31) had an adequate depth of coverage (>1000 reads per nucleotide) for haplotype estimation with Nano-Q. The median number of haplotypes identified per patient was 9.5 (IQR: 4.0 - 15.8). The total number of haplotypes generated for all patients with DENV1 (n=8), DENV2 (n=51) and DENV3 (n=31) serotypes were 40, 541, and 376 respectively.

To assess the Shannon entropy (SE) of DENV genomes at consensus level only DENV2 and DENV3 alignments were considered as the DENV1 alignment had only 8 sequences. The average SE across the genome was statistically significantly higher for DENV2 compared to DENV3 [0.0041 (SD: 0.037) vs. 0.0020 (SD 0.0305), p < 0.0001, Mann Whitney U test] (Figure 3A, 3B). A total of 69 and 21 sites had significantly higher SE (> mean + 2SD) in DENV2 and DENV3 alignments respectively (Table 1). One of these positions in DENV2 (position 1129, NS2A gene) was identified as undergoing diversifying selection by the FUBAR algorithm. The highest average of sites with significantly high SE was seen in NS2A gene when normalised for the gene length in both serotypes.

**Figure 3.**
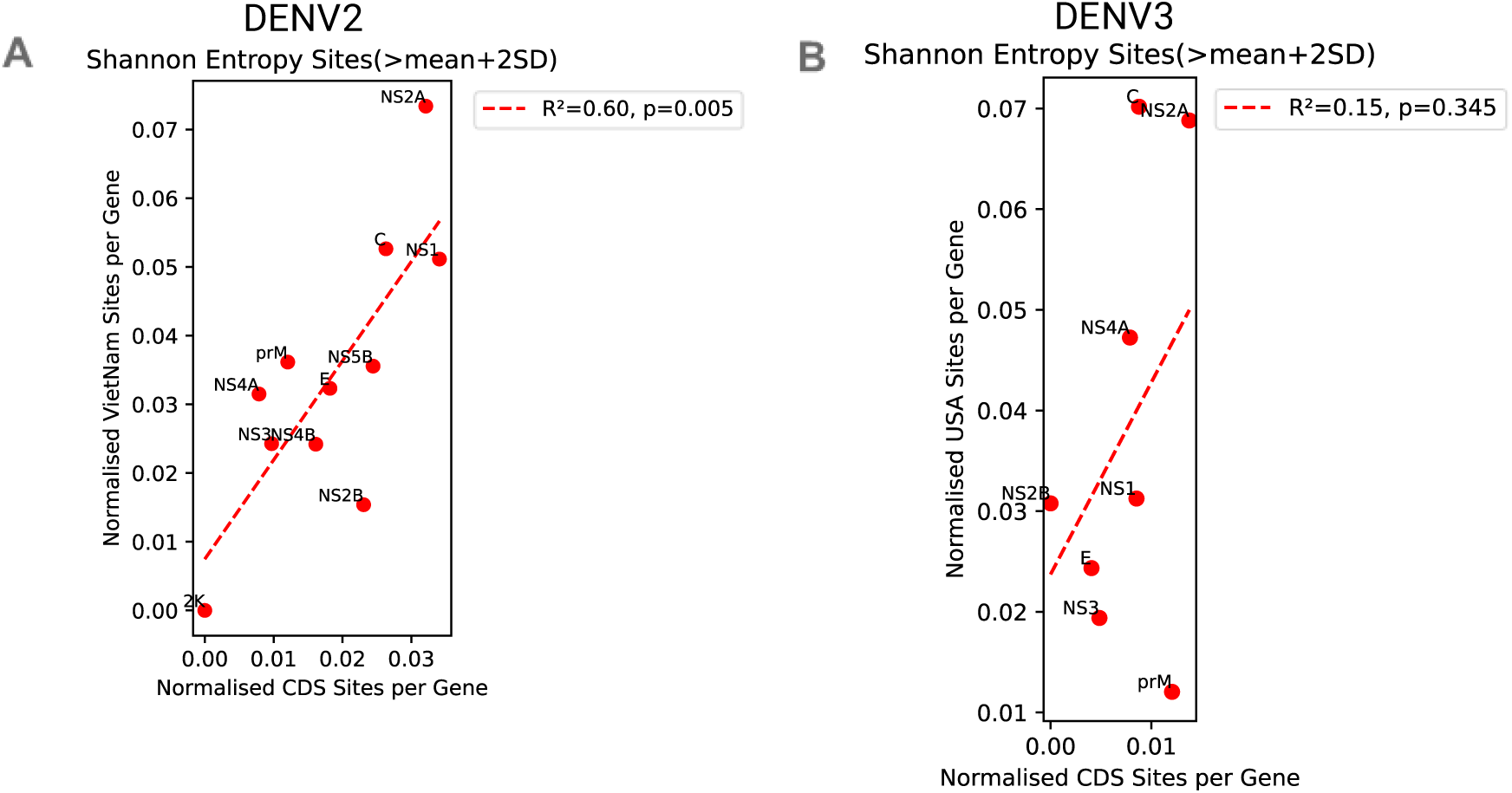
Scatter plots of the normalised sites of entropy and normalised sites of selection. (A) Scatter plot of high entropy sites between DENV2 CDS sequences and the DENV2 sequences from the Vietnam cohort (R^2^=0.60, p=0.05). (B) Scatter plot of high entropy sites between DENV3 CDS sequences and the DENV2 sequences from the Florida, USA cohort (R^2^=0.15, p=0.345).

**Table 1.** Summary of number of sites with Shannon Entropy > mean + 2SD for each gene in DENV2 and DENV3 alignments of CDS consensus sequences

| **Gene** | **DENV2** |  | **DENV3** |  |
| --- | --- | --- | --- | --- |
|  | Sites with<br>SE>mean+2SD | Normalized<br>value* | Sites with<br>SE>mean+2SD | Normalized<br>value* |
| **C** | 3 | 0.026 | 1 | 0.009 |
| **prM** | 2 | 0.012 | 2 | 0.012 |
| **E** | 9 | 0.018 | 2 | 0.004 |
| **NS1** | 2 | 0.006 | 3 | 0.009 |
| **NS2A** | 7 | 0.032 | 3 | 0.014 |
| **NS2B** | 3 | 0.023 | 0 | 0 |
| **NS3** | 6 | 0.01 | 3 | 0.005 |
| **NS4A** | 1 | 0.008 | 1 | 0.008 |
| **2K** | 0 | 0 | 0 | 0 |
| **NS4B** | 4 | 0.016 | 0 | 0 |
| **NS5B** | 22 | 0.024 | 6 | 0.007 |

(\*normalized for the length of each gene)

For external validation, publicly available DENV1 and DENV2 sequences (397 and 125 respectively) form an outbreak in Vietnam (2006-2008) and 173 DENV3 sequences from Florida, USA (2023-2024) were analysed. The average SE estimates were 0.0585 (SD= 0.1811), 0.0605 (SD=0.1706), 0.0391 (SD= 0.0956) for DENV1, DENV2, and DENV3 respectively (Supplementary Figure 1A, 1B and 1C) and the difference between DENV2 and DENV3 was statistically significant (p <0.0001, Mann Whitney U test). In these alignments 84 (DENV1), 65 (DENV2), and 51 (DENV3) sites had significantly higher SE values (> mean + 2SD). The average number of such sites normalised by gene length is shown in Supplementary Table 1, which confirmed that NS2A had the highest value in all three serotypes except that the capsid gene also tied for the top position in DENV3. The number of sites with high SE per gene were similarly distributed between the DENV2 sequences of CDS and the corresponding external dataset (R^2^ = 0.6, p=0.005), and less so for DENV3 (R^2^ = 0.15, p=0.345). Up to 7 (DENV1), 8 (DENV2) and 4 (DENV3) codons were identified as undergoing diversifying selection by at least one algorithm in the external datasets, but only 1 site (position 3326 in NS5B of DENV1) was confirmed by all algorithms (Supplementary Table 2 and 3).

For haplotype level analysis of mutability, all three serotypes including DENV1 was considered. The average SE in haplotype alignments were 0.00146 (SD= 0.0152), 0.00123 (SD= 0.0051) and 0.00189 (SD= 0.0052) in DENV1, DENV2 and DENV3 respectively, and this difference was statistically significant (p<0.0001, Kruskal-Wallis test). A total of 28 (DENV1), 41 (DENV2) and 89 (DENV3) sites had a significantly higher SE. When normalised for length of each gene, NS1 had the highest average for sites with high SE in both DENV1 and DENV2 alignments while capsid, NS1, and NS2B genes all had high values in DENV3 alignment (Table 2). Also, 15 (DENV1), 43 (DENV2), and 47 (DENV3) codons were under diversifying selection as identified by at least one of the algorithms, with 3, 14, and 11 codons respectively found to be under diversifying selection by all three algorithms (Table 3). A descriptive account of diversifying selection sites identified by each of the algorithms for each serotype can be found in supplementary table 4. An external validation could not be performed at the haplotype level as equivalent data were not available in the public domain.

**Table 2.** Summary of sites with a Shannon entropy > mean + 2SD normalised for each gene for DENV1 DENV2 and DENV3 haplotype sequences.

|  | DENV1 |  | DENV2 |  | DENV3 |  |
| --- | --- | --- | --- | --- | --- | --- |
| GENE | Number of sites | Normalised value* | Number of sites | Normalised value* | Number of sites | Normalised value* |
| **C** | 2 | 1.7544 | 1 | 0.8772 | 9 | 7.8947 |
| **PRM** | 1 | 0.6024 | 3 | 1.8072 | 5 | 3.0120 |
| **E** | 3 | 0.6061 | 10 | 2.0284 | 19 | 3.8540 |
| **NS1** | 11 | 3.1250 | 13 | 3.6932 | 19 | 5.3977 |
| **NS2A** | 4 | 1.8349 | 5 | 2.2936 | 6 | 2.7523 |
| **NS2B** | 1 | 0.7692 | 1 | 0.7692 | 9 | 6.9231 |
| **NS3** | 6 | 0.9693 | 8 | 1.2924 | 22 | 3.5541 |

(\*normalised for the length of each gene)

**Table 3.** Summary of identified sites of diversifying selection from all the algorithms and any of the algorithms FEL FUBAR and MEME the haplotype sequences for each gene for each serotype.

| DENV1 |  |  |  |  |
| --- | --- | --- | --- | --- |
| Gene | Number of Sites Identified by all 3 algorithms | Number of Sites Identified by all 3 algorithms normalised* | Number of Sites identified by any of the algorithms | Number of Sites identified by any of the algorithms normalised* |
| *C* | 0 | 0 | 1 | 0.877 |
| *prM* | 1 | 0.602 | 1 | 0.602 |
| *E* | 0 | 0 | 2 | 0.404 |
| *NS1* | 0 | 0 | 4 | 1.136 |
| *NS2A* | 0 | 0 | 1 | 0.459 |
| *NS2B* | 0 | 0 | 0 | 0 |
| *NS3* | 2 | 0.323 | 6 | 0.969 |

*DENV2*
| *Gene* | Number of Sites Identified by all 3 algorithms | Number of Sites Identified by all 3 algorithms normalised* | Number of Sites identified by any of the algorithms | Number of Sites identified by any of the algorithms normalised* |
| --- | --- | --- | --- | --- |
| *C* | 0 | 0 | 5 | 4.386 |
| *prM* | 0 | 0 | 6 | 3.614 |
| *E* | 3 | 0.609 | 10 | 2.028 |
| *NS1* | 3 | 0.852 | 8 | 2.273 |
| *NS2A* | 3 | 1.376 | 5 | 2.294 |
| *NS2B* | 1 | 0.769 | 1 | 0.769 |
| *NS3* | 4 | 0.646 | 8 | 1.292 |

*DENV3*
| *Gene* | Number of Sites Identified by all 3 algorithms | Number of Sites Identified by all 3 algorithms normalised* | Number of Sites identified by any of the algorithms | Number of Sites identified by any of the algorithms normalised* |
| --- | --- | --- | --- | --- |
| *C* | 2 | 1.754 | 9 | 7.895 |
| *prM* | 2 | 1.205 | 8 | 4.819 |
| *E* | 1 | 0.203 | 10 | 2.028 |
| *NS1* | 2 | 0.568 | 5 | 1.420 |
| *NS2A* | 0 | 0 | 2 | 0.917 |
| *NS2B* | 3 | 2.308 | 4 | 3.077 |
| *NS3* | 1 | 0.162 | 9 | 1.454 |

(\*normalised for the length of each gene)

### 3.1 Mutation hotspots

Mutation hotspots were defined as sites with a SE > mean +2SD, contained within a codon undergoing diversifying selection as detected by at least one selection algorithm (Figure 4A,4B and 4C). At the consensus level, only 1 site in DENV2 (Position 1129 in the NS2A gene) and 0 sites in DENV3 alignment met this criterion (DENV1 alignment was not considered) whereas in the external datasets 2 (DENV1), 2 (DENV2), 3 (DENV3) sites met the same criteria (Supplementary Table 5), without an overlap with the CDS dataset with regard to specific positions. At the haplotype level, more hotspots were identified (DENV1: 15, DENV2: 23, DENV3: 17) (supplementary table 6) and of these, 3 (DENV1), 13 (DENV2) and 8 (DENV3) codons were recognised by all three algorithms as undergoing diversifying selection (Table 4).

**Figure 4.**
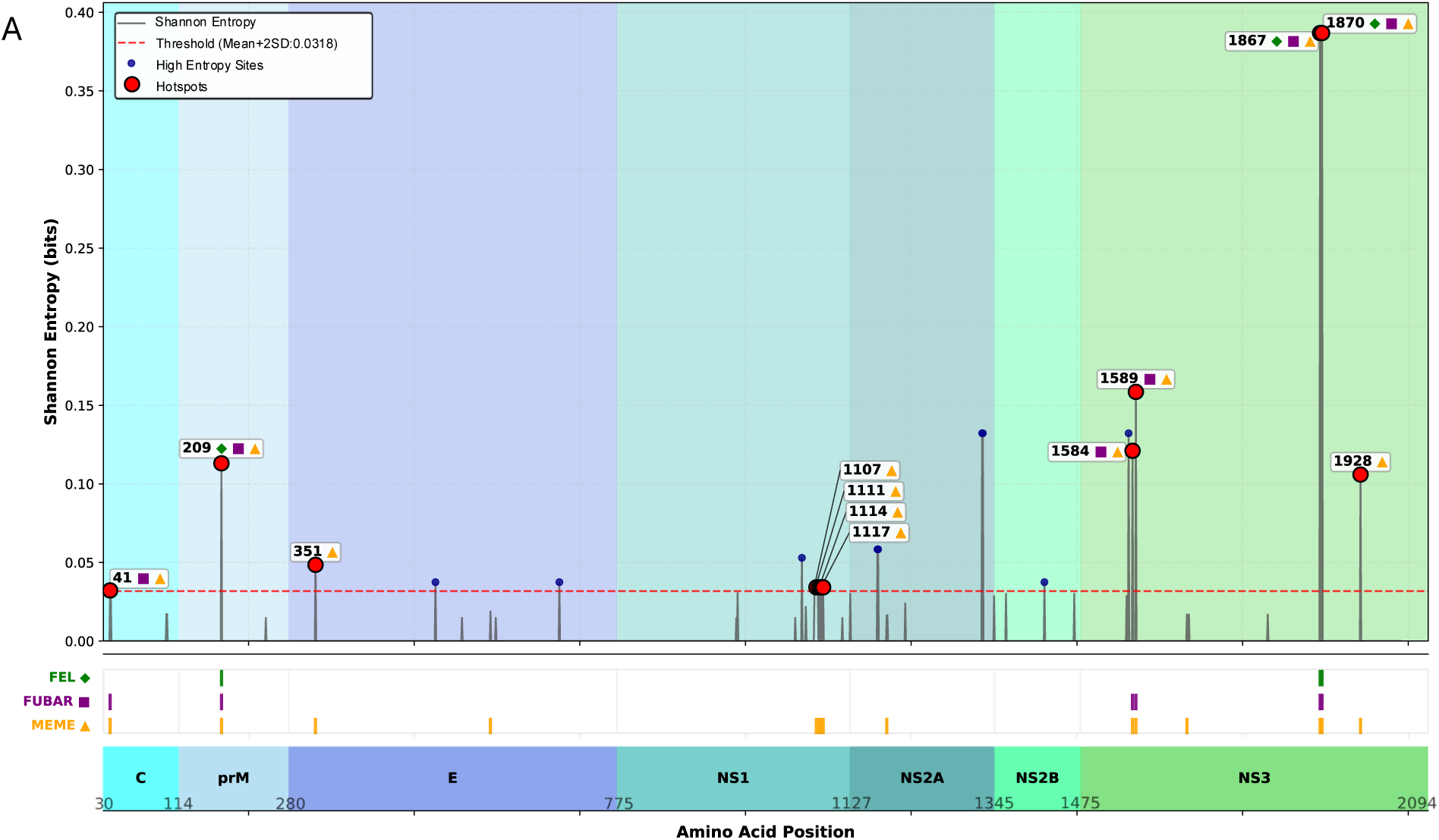
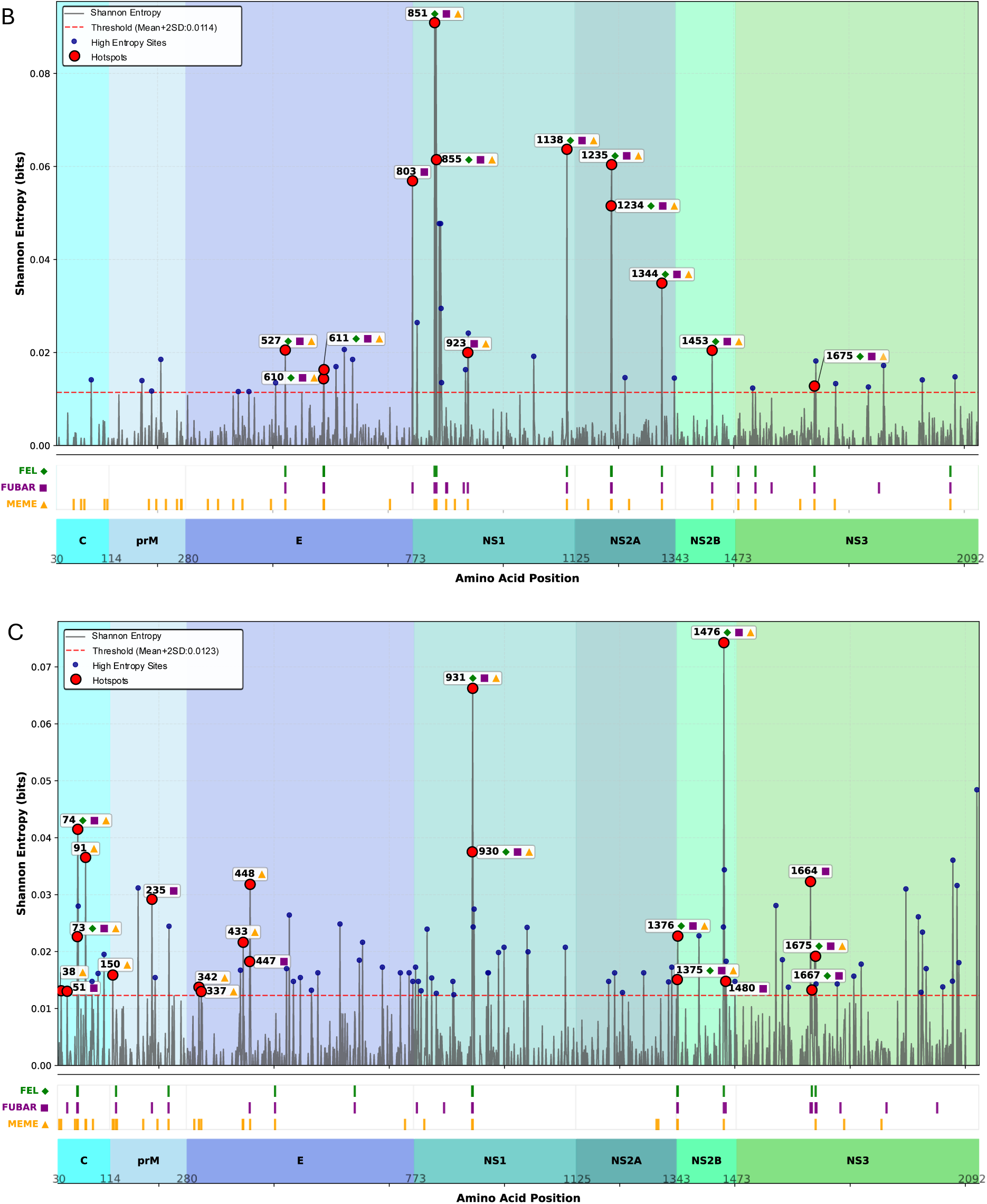
Graphs of Shannon entropy values against the amino acid position for (A) DENV1 (B)DENV2 (C)DENV3 haplotype sequences. The red dotted line marks Shannon entropy cutoff that was set at mean+2SD. Each blue point indicates the sites that have entropy values over the threshold. The sites of diversifying selection are marked with green, purple or yellow to indicate FEL FUBAR or MEME algorithm respectively. Hotspots are marked with red points and labelled with the amino acid position and marked with green diamonds for FEL, purple squares for FUBAR and yellow triangles for MEME to indicate the selection algorithm. The amino acid coordinates for the relevant viral genes are also marked on the x-axis.

**Table 4.**
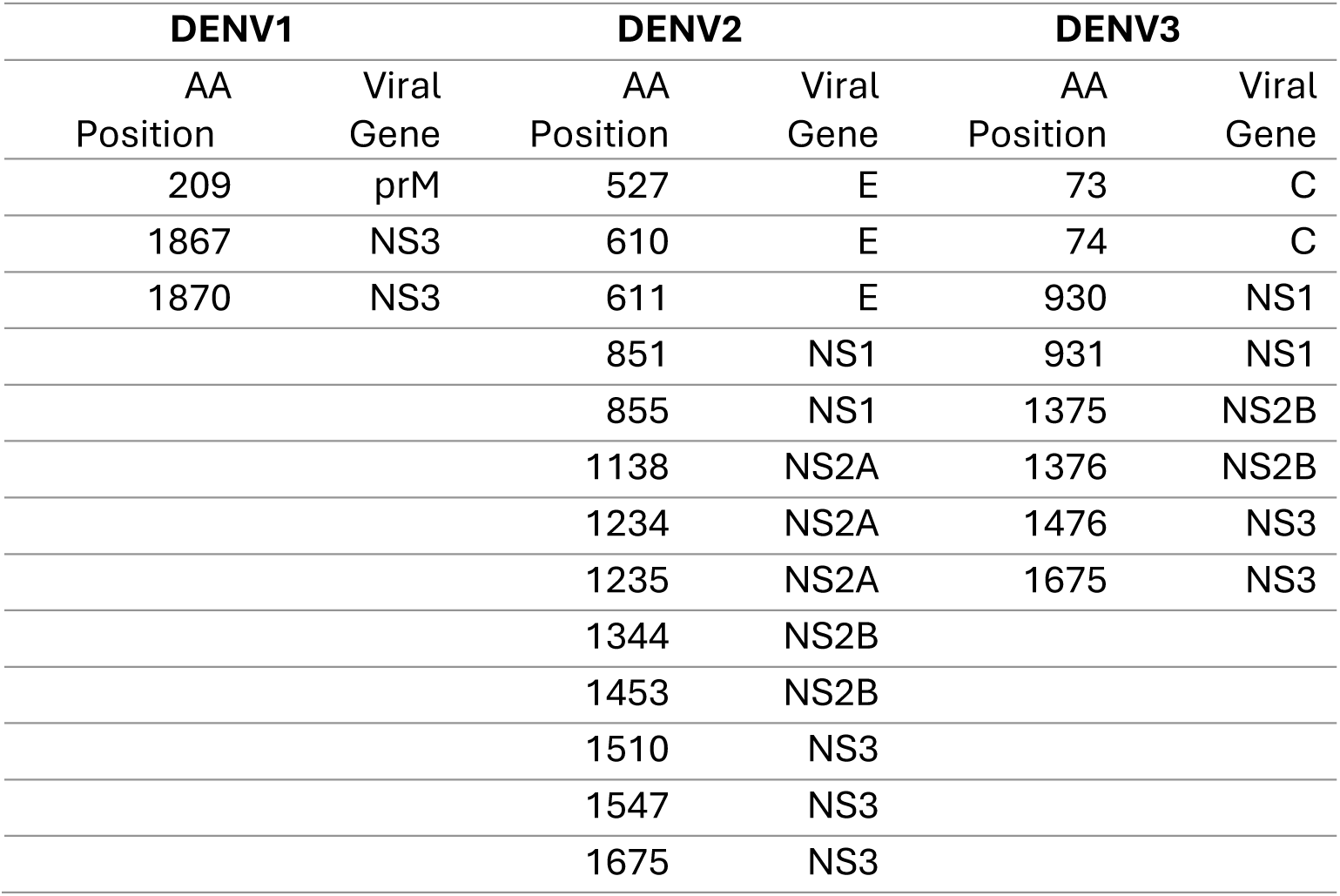
Hotspots as identified by all 3 selection algorithms (FEL FUBAR and MEME) and Shannon entropy (sites over mean+2SD) for DENV1 DENV2 and DENV3 and the corresponding viral genes. All hotspots identified for DENV1 DENV2 and DENV3 can be found in the supplementary table 6.

| DENV1 |  | DENV2 |  | DENV3 |  |
| --- | --- | --- | --- | --- | --- |
| AA | Viral Gene | AA | Viral Gene | AA | Viral Gene |
| Position |  | Position |  | Position |  |
| 209 | prM | 527 | E | 73 | C |
| 1867 | NS3 | 610 | E | 74 | C |
| 1870 | NS3 | 611 | E | 930 | NS1 |
|  |  | 851 | NS1 | 931 | NS1 |
|  |  | 855 | NS1 | 1375 | NS2B |
|  |  | 1138 | NS2A | 1376 | NS2B |
|  |  | 1234 | NS2A | 1476 | NS3 |
|  |  | 1235 | NS2A | 1675 | NS3 |
|  |  | 1344 | NS2B |  |  |
|  |  | 1453 | NS2B |  |  |
|  |  | 1510 | NS3 |  |  |
|  |  | 1547 | NS3 |  |  |
|  |  | 1675 | NS3 |  |  |

## 4. Discussion

This study explored the mutability of the DENV genome at both consensus and haplotype levels using viral sequences isolated from Sri Lanka between 2017 – 2020, dominated by serotypes 2 and 3. At the consensus level DENV2 sequences had a higher average SE across the genome compared to DENV3, but this pattern was reversed when haplotypes were compared. An external validation which could only be done at consensus level confirmed these findings. At the consensus level the largest number of highly mutable sites were seen in NS2A gene when adjusted for gene length in all serotypes, while at the haplotype level the NS1 gene had the most mutable sites in DENV1 and DENV2.

Unlike HIV or HCV infections, dengue is a short-lived infection in humans without chronic viraemia. DENV is invariably cleared by immunocompetent surviving hosts, typically within 4-5 days from the onset of fever. Given the limited time to mutate within an individual host, and as confirmed by our previous work comparing both dengue and HCV, the former shows comparatively far fewer mutations both at consensus and haplotype levels (T. N. Adikari et al., 2020; Riaz et al., 2021; Riaz et al., 2022). Yet even for a universally cleared infection like dengue, if one serotype has a higher mutability than others, it may still survive longer against the host immune system, with subtle differences in the externally manifest disease phenotype. Phylogenetically, DENV1 and DENV3 are more closely related to each other while DENV2 is more distant and distinct(“Real-time tracking of dengue virus evolution,” 2025). Meta-analyses on serotype-specific differences on dengue clinical illness have found that DENV2 infections are associated with more cases of severe dengue in children (Sangkaew et al., 2021)), higher overall mortality (Guo et al., 2017) and higher risk of dengue shock syndrome (Huy et al., 2013). While our study cannot directly link high mutability of DENV2 to such previously described clinical associations, it poses an interesting hypothesis that warrants further investigation. The NS2A gene consistently had more sites with higher SE when adjusted for gene length in both CDS and external datasets, regardless of the serotype when considered at the consensus level. This gene codes for a 22kDa hydrophobic protein which is critical for dengue RNA synthesis, virion assembly and for countering the host innate immune response by inhibiting interferon signalling (Muñoz-Jordan, Sánchez-Burgos, Laurent-Rolle, & García-Sastre, 2003)). Structurally, the NS2 protein gets integrated to the host cell endoplasmic reticulum (ER) with its N-terminus within the ER complex and its C-terminus projecting into the cytosol, with five transmembrane domains embedded into the ER lipid bilayer (Xie, Gayen, Kang, Yuan, & Shi, 2013). Apart from NS2A gene, both capsid and NS1 genes also had more highly mutable sites in both DENV2 and DENV3 sequences from the CDS cohort. Capsid is critical for the structural integrity of the virus, while NS1 is a non-structural protein critical for viral replication (Avirutnan et al., 2006; Lindenbach & Rice, 1997) and initiation of plasma leakage which is a preceding event for most clinical complications in dengue. Overall, there was good concordance between CDS and the external dataset of DENV2 sequences regarding the number of highly mutable sites per gene, but less so for DENV3. This could have been influenced by the smaller alignment size for DENV3 sequences in CDS resulting in less available information for analysis. Also, since the external dataset from Florida, USA was not linked to a publication, it was not possible to deduce if all sequences were in fact linked to the same outbreak. Nevertheless, because these sequences were isolated between adjacent calendar years (2023 – 2024) we assumed the sequences were from the same outbreak, although it is likely that at least a small subset may be imported infections from other countries or other states and therefore linked to different outbreaks.

The haplotype level analysis which could only be done for CDS sequences showed a contrasting pattern of higher average SE in DENV3 alignment compared to DENV2. We previously showed that in the latter part of the sampling period of CDS (in 2019), a major DENV3 dominated outbreak replaced the existing DENV2 dominated outbreak in the community(Maduranga et al., 2023). Such a serotype shift occurs when a new fit variant from a different serotype is introduced to a community with waning or non-existent immunity to that serotype. All four serotypes are known to be endemic in Sri Lanka, but from time-to-time disproportionate outbreaks associated with serotype switches have occurred (e.g., DENV1 in 2009, DENV2 in 2016/17 and DENV3 in 2019), highlighting the waning serotype specific immunity over time. Thus, a DENV3 infection in a host with DENV2 directed immunity (or no dengue-specific immunity), may allow the virus to survive longer by mutating faster before eventually being wiped out. Such changes may only be apparent at haplotype level because the consensus level analyses are biased towards the high abundance haplotypes ignoring the minor variant quasi-species. This example highlights how additional information on viral evolution can be elicited by studying haplotypes.

Some sites with high SE were also identified as sites with diversifying selection by at least one of the three algorithms used for this purpose. Such mutation hotspots were spotted more at the haplotype level analyses than at the consensus level with no overlap between results of each analysis type (haplotype vs. consensus). The hotspots identified at the consensus level are more informative of virus evolution in the community (e.g., predicting rising or waning intensity of an outbreak), whereas the hotspots identified at the haplotype level may be more relevant to understand which parts of the virus genome are mutation-prone. The position of each mutation hotspot identified, and the known functional significance of the peptide subunit they are contained within is summarised in supplementary table 7 for each serotype. None of the patients whose samples were sequenced for this project had severe dengue and were none were admitted to an intensive care unit or died. Therefore, it was not possible to associate haplotype level mutation hotspots to infection outcomes in this study, but the feasibility of such analyses is evident if applied to patient populations with more diverse clinical outcomes.

This study had several limitations. Firstly, the consensus level analysis was restricted to DENV2 and DENV3 sequences with inequal sample sizes. Sequence availability was determined by the dengue outbreaks in the community at the time of sampling. Secondly, not all available viraemic samples could be sequenced to an adequate depth to generate haplotypes. As this cohort was hospital-based, most eligible people were recruited on days 3 and 4 of fever after being referred by a primary care physician or from the out-patients department of the hospital for persistent fever. Thus, the level of viraemia was most likely on the decline for most patients. Thirdly, only one sample was available per patient, making it impossible to observe longitudinal changes in haplotype diversity. However, it should be noted that given that dengue viraemia is very short lasting (4-5 days), significant changes in haplotypes are unlikely to be observed even with daily sampling. Fourthly, lack of publicly available near-full-length DENV sequences linked to meta-data (year and country of isolation) and a publication to explain the context of sampling limited our capacity to identify suitable datasets for external validation, especially for DENV3. Although more DENV3 genomes were available from Nicaragua and Cuba, the sequence quality was less optimal (long stretches of ambiguous nucleotides or gaps) compared to the selected set of sequences from USA. As no DENV4 sequences were identified in CDS, this serotype was not considered in this paper. Finally, despite inputting equimolar concentrations of DNA (which was reverse transcribed from RNA), the sequencing depth achieved per sample was highly variable. This creates a bias as samples sequenced to a higher depth had more information available to potentially discover more minor variants(T. N. Adikari et al., 2020). To limit this effect and false discovery, only haplotypes occurring at a frequency > 0.1% were considered. Another problem created by the uneven sequencing output is that to maximize the number of samples and sequences the length of the haplotype reconstructed was limited to 6000 nucleotides which meant that the NS4A, NS4B and NS5 genes had to be omitted from the haplotype-level analysis.

## 5. Conclusions

This in-depth analysis of the mutability of the dengue virus genome using near-full-length sequences from two continuous DENV2 and DENV3 dominated outbreaks in Sri Lanka revealed consistent, but different patterns of serotype-specific mutation propensities at consensus and haplotype level analyses. Haplotypes showed more mutations and codons under diversifying selection than those visible at consensus level. Advances in RNA virus near-full-length genome amplification assays, introduction of long-read third generation sequencing platforms and new bioinformatic tools to reconstruct viral haplotypes have made it easier to study viral mutations at haplotype level.

## 7. Supporting Statements

### 7.1 Ethics approvals

The work described in this paper is covered by human ethics committee approvals from the University of Colombo, Sri Lanka (EC-17-080) and The University of New South Wales, Australia (HC220706). All patients provided informed written consent prior to recruitment and sample collection.

## Acknowledgements

We would like to acknowledge Tonia Russell who assisted with some of the sequencing runs.

## 7.2 Data Availability

The raw sequencing data will be uploaded to Genbank and made publicly available. The consensus sequences of the CDS cohort have been uploaded to Genbank previously (OR393874-OR394054).

## 7.3 Declarations

I.W.D. manages a fee-for-service sequencing facility at the Garvan Institute and is a customer of Oxford Nanopore Technologies and Pacific BioSciences but has no further financial relationship. I.W.D. has received travel and accommodation expenses from Oxford Nanopore Technologies. The authors declare no other competing financial or nonfinancial interests.

## 8. Supplementary figures and tables

**Supplementary table 1:**
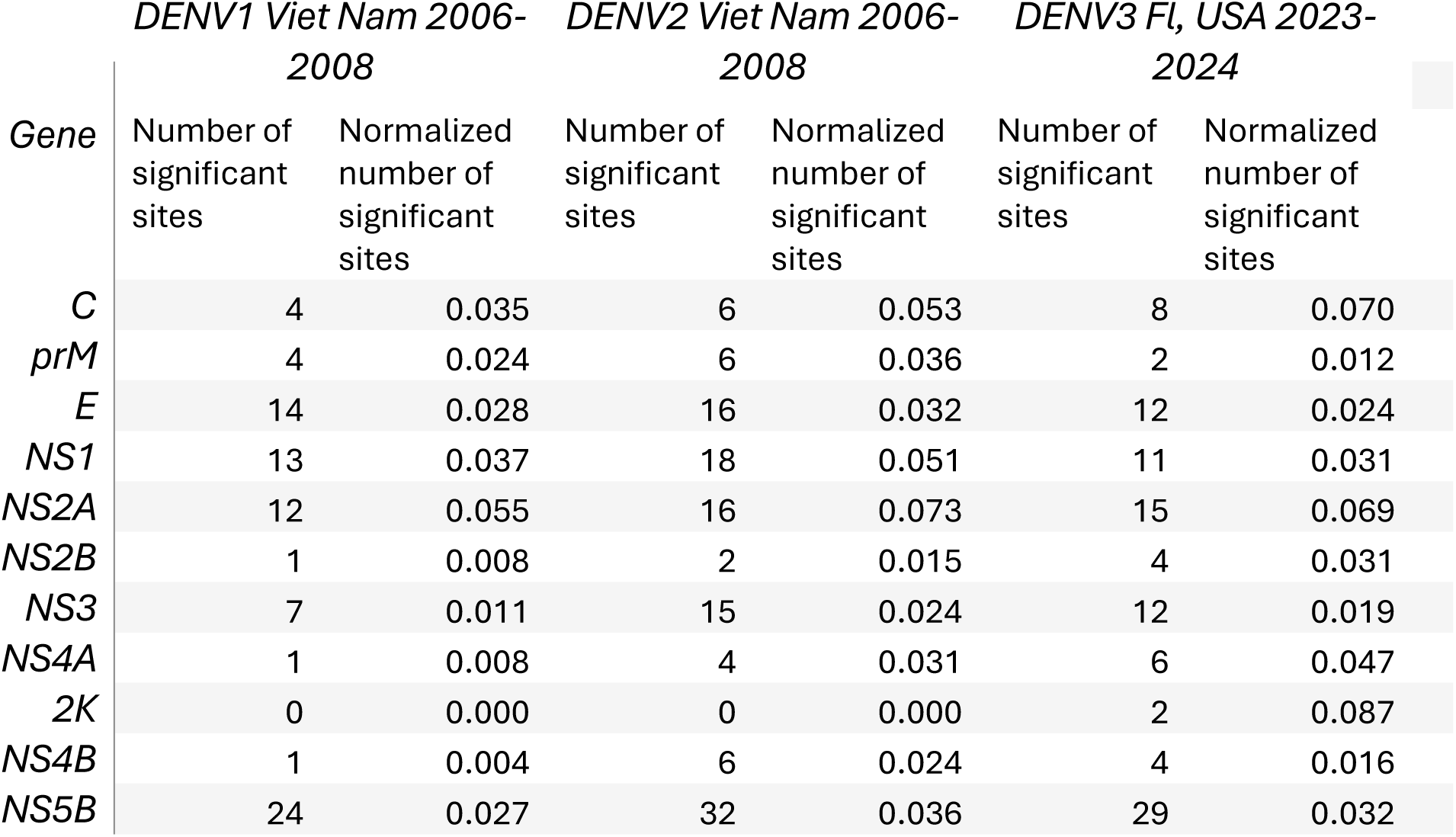
Number of sites with Shannon entropy > 2SD + mean per gene and normalized number of such sites for each gene for DENV1 and DENV2 sequences from Vietnam and DENV3 sequences from Florida, USA (the datasets used for external validation of consensus level results)

| *Gene* | *DENV1 Viet Nam 2006-2008* |  | *DENV2 Viet Nam 2006-2008* |  | *DENV3 Fl, USA 2023-2024* |  |
| --- | --- | --- | --- | --- | --- | --- |
|  | Number of significant sites | Normalized number of significant sites | Number of significant sites | Normalized number of significant sites | Number of significant sites | Normalized number of significant sites |
| *C* | 4 | 0.035 | 6 | 0.053 | 8 | 0.070 |
| *prM* | 4 | 0.024 | 6 | 0.036 | 2 | 0.012 |
| *E* | 14 | 0.028 | 16 | 0.032 | 12 | 0.024 |
| *NS1* | 13 | 0.037 | 18 | 0.051 | 11 | 0.031 |
| *NS2A* | 12 | 0.055 | 16 | 0.073 | 15 | 0.069 |
| *NS2B* | 1 | 0.008 | 2 | 0.015 | 4 | 0.031 |
| *NS3* | 7 | 0.011 | 15 | 0.024 | 12 | 0.019 |
| *NS4A* | 1 | 0.008 | 4 | 0.031 | 6 | 0.047 |
| *2K* | 0 | 0.000 | 0 | 0.000 | 2 | 0.087 |
| *NS4B* | 1 | 0.004 | 6 | 0.024 | 4 | 0.016 |
| *NS5B* | 24 | 0.027 | 32 | 0.036 | 29 | 0.032 |

**Supplementary table 2.**
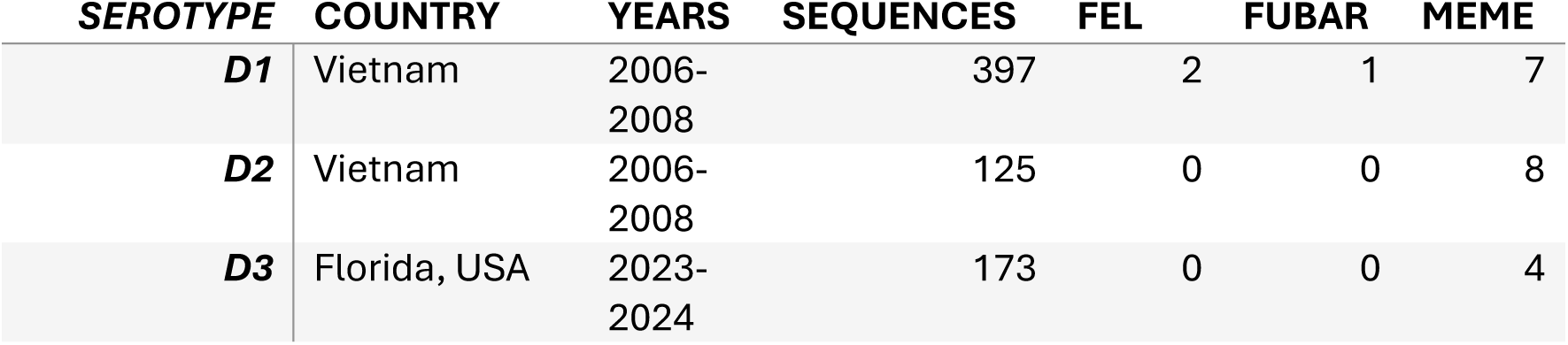
Summary of external datasets and sites of diversifying selection identified by each algorithm

**Supplementary table 3.**
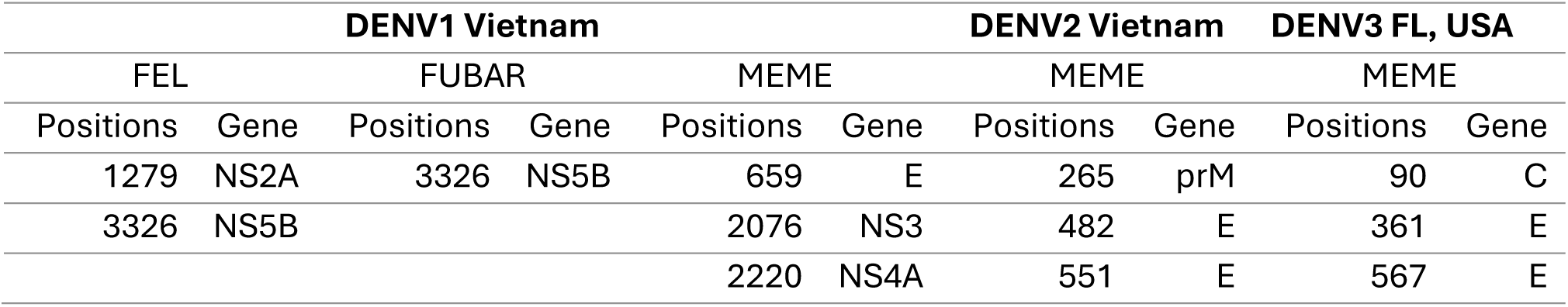
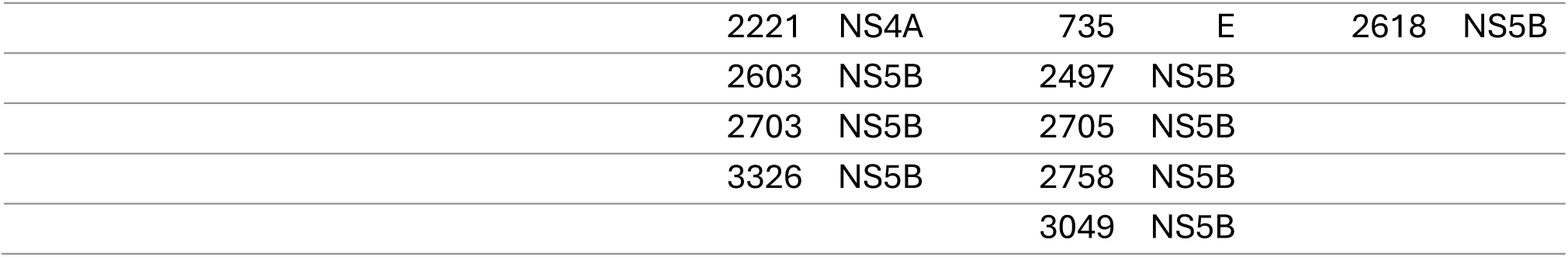
Sites with diversifying selection identified by each algorithm per serotype in external validation datasets

**Supplementary table 4:**
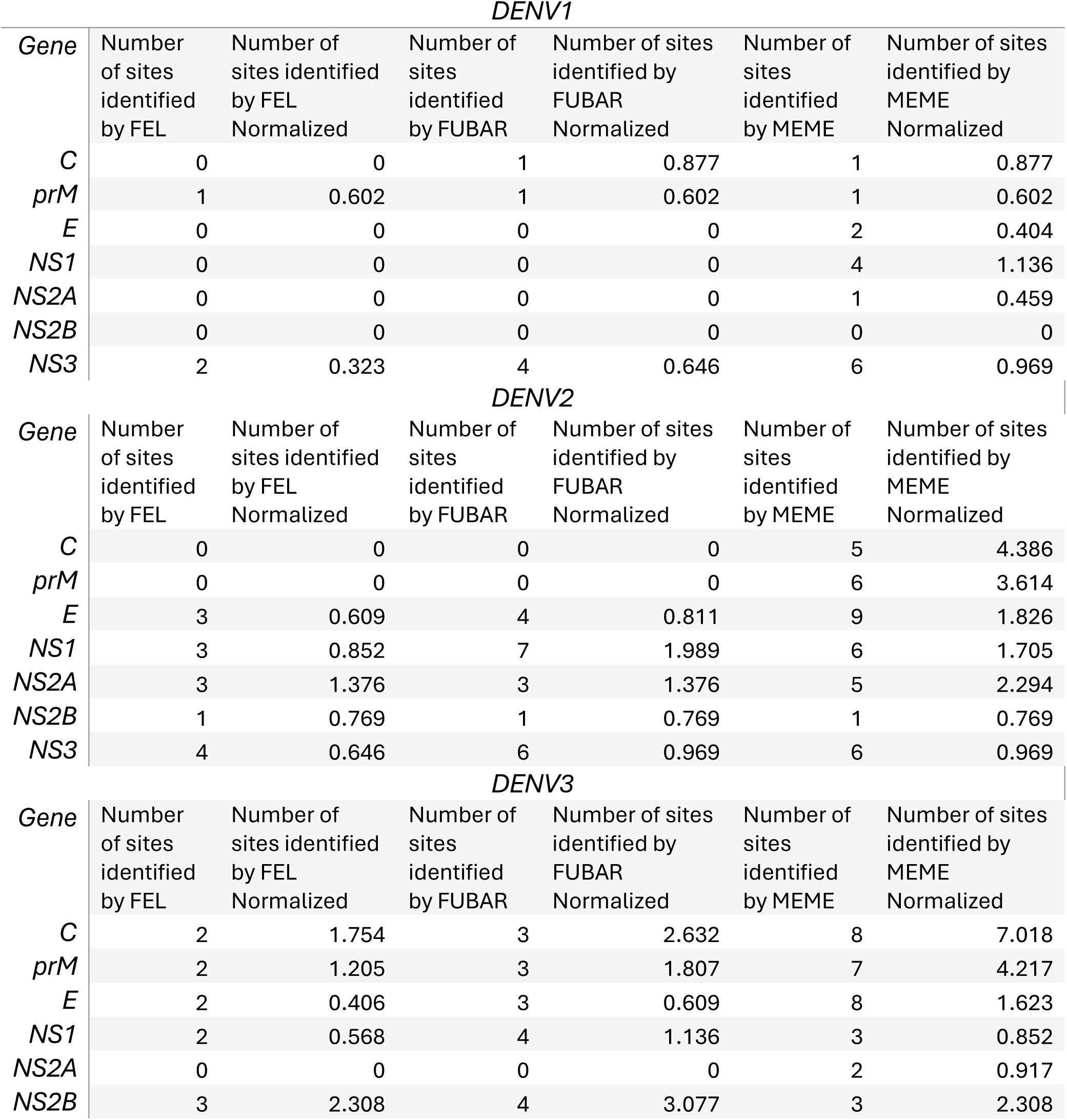
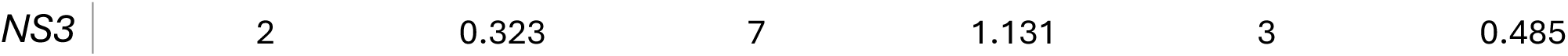
Summary of sites undergoing diversifying selection identified by each of the algorithms FEL. FUBAR, MEME in haplotype alignments in the CDS cohort for each of the serotypes, stratified by their location in the genome.

**Supplementary table 5:** Mutation hotspots identified from each of the external datasets for each serotype

| DENV1 Vietnam |  | DENV2 Vietnam |  | DENV3 FL, USA |  |
| --- | --- | --- | --- | --- | --- |
| **659** | E | 2758 | NS5B | 90 | C |
| **3326** | NS5B | 3049 | NS5B | 361 | E |
|  |  |  |  | 2618 | NS5B |

**Supplementary table 6:**
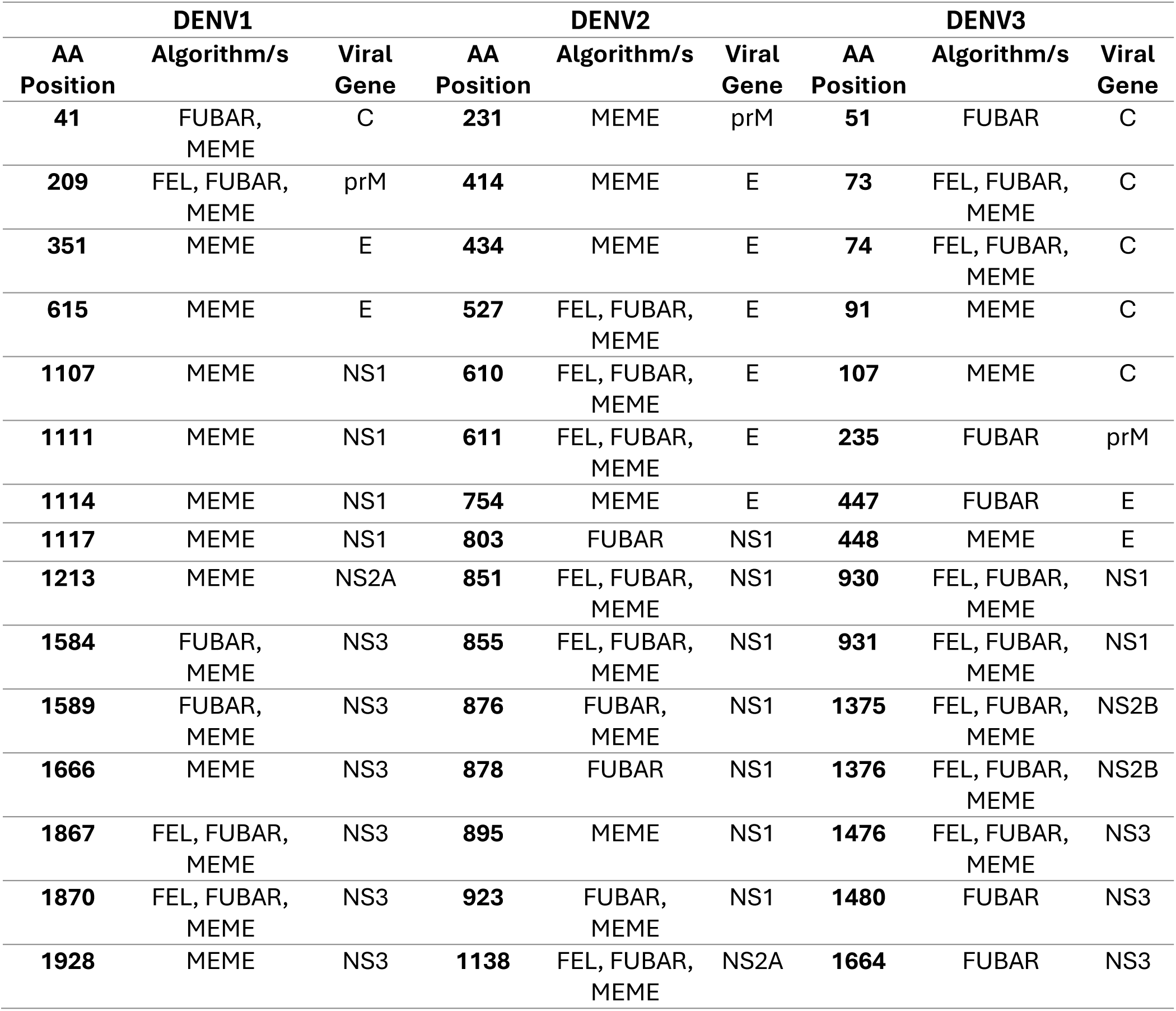
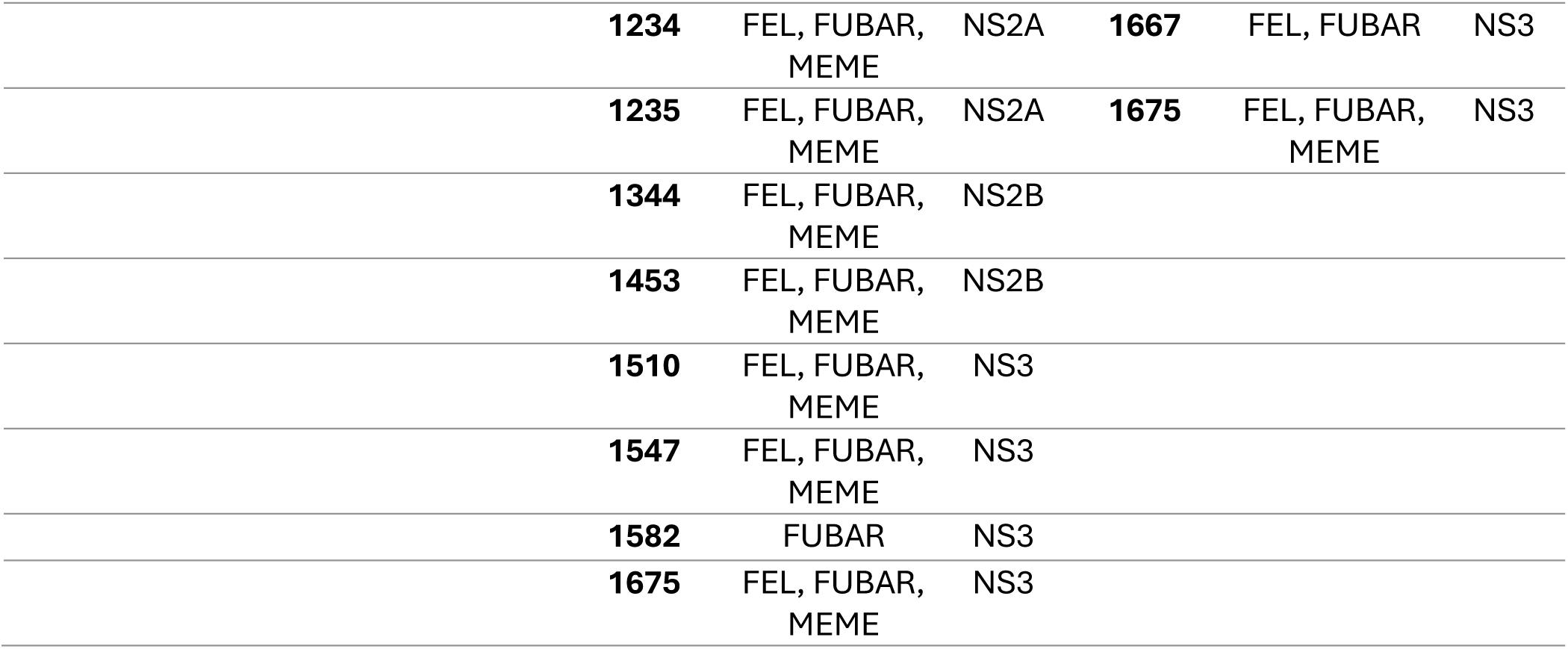
Summary of all the hotspots identified at the haplotype level for the CDS cohort with the supporting selection algorithms.

**Supplementary table 7:** The mutation hotspots identified at the haplotype level and the structural elements that they correspond to within the DENV genome.

| *Mutation Hotspot* | *Gene* | *Gene Position* | *Structural Element* | *PDB_IDs* | *Functional Note* |
| --- | --- | --- | --- | --- | --- |
| **DENV1** |  |  |  |  |  |
| 41 | C | 11 | N-terminal disordered region | 1R6R |  |
| 209 | prM | 65 | pr domain (chaperone function) | 3C5X |  |
| 351 | E | 41 | Domain I - central | 4GT0, 4CML, 4B03 |  |
| 1107 | NS1 | 302 | $\hat{I}^2$-ladder domain | 4O6B, 5IY3 | $\hat{I}^2$-ladder domain involved in NS1 dimerization and immune system interaction |
| 1111 | NS1 | 306 | $\hat{I}^2$-ladder domain | 4O6B, 5IY3 | $\hat{I}^2$-ladder domain involved in NS1 dimerization and immune system interaction |
| 1114 | NS1 | 309 | $\hat{I}^2$-ladder domain | 4O6B, 5IY3 | $\hat{I}^2$-ladder domain involved in NS1 dimerization and immune system interaction |
| 1117 | NS1 | 312 | $\hat{I}^2$-ladder domain | 4O6B, 5IY3 | $\hat{I}^2$-ladder domain involved in NS1 dimerization and immune system interaction |
| 1584 | NS3 | 79 | Protease domain | 2VBC, 2JLQ, 5JRZ | Protease domain cleaves the viral polyprotein |
| 1589 | NS3 | 84 | Protease domain | 2VBC, 2JLQ, 5JRZ | Protease domain cleaves the viral polyprotein |
| 1867 | NS3 | 362 | Helicase domain II | 2VBC, 2JLQ, 5JRZ | Helicase domain unwinds RNA during replication |
| 1870 | NS3 | 365 | Helicase domain II | 2VBC, 2JLQ, 5JRZ | Helicase domain unwinds RNA during replication |
| 1928 | NS3 | 423 | Helicase domain II | 2VBC, 2JLQ, 5JRZ | Helicase domain unwinds RNA during replication |
| **DENV2** |  |  |  |  |  |
| 527 | E | 217 | Domain II - fusion loop | 1OAN, 1TG8, 2JSF | Fusion loop mediates membrane fusion during viral entry |
| 610 | E | 300 | Domain III - receptor binding | 1OAN, 1TG8, 2JSF | Domain III is involved in receptor binding and is a major target for neutralizing antibodies |
| 611 | E | 301 | Domain III - receptor binding | 1OAN, 1TG8, 2JSF | Domain III is involved in receptor binding and is a major target for neutralizing antibodies |
| 803 | E | 493 | Transmembrane region | 1OAN, 1TG8, 2JSF |  |
| 851 | NS1 | 48 | Wing domain | 4O6B, 5K6K |  |
| 855 | NS1 | 52 | Wing domain | 4O6B, 5K6K |  |
| 923 | NS1 | 120 | Wing domain | 4O6B, 5K6K |  |
| 1138 | NS1 | 335 | $\hat{I}^2$-ladder domain | 4O6B, 5K6K | $\hat{I}^2$-ladder domain involved in NS1 dimerization and immune system interaction |
| 1234 | NS2A | 79 | Central transmembrane region | 2KQI |  |
| 1235 | NS2A | 80 | Central transmembrane region | 2KQI |  |
| 1344 | NS2A | 189 | C-terminal domain | 2KQI |  |
| 1453 | NS2B | 80 | NS3 protease cofactor region | 2M9P, 2FOM | NS2B cofactor is essential for NS3 protease activity |
| 1675 | NS3 | 172 | Linker region | 2VBC, 2WHX, 2JLQ |  |
| **DENV3** |  |  |  |  |  |
| 38 | C | 8 | N-terminal disordered region | 1R6R |  |
| 51 | C | 21 | $\hat{I}^{\pm 1}$ helix (RNA binding) | 1R6R | RNA binding region packages viral genome |
| 73 | C | 43 | $\hat{I}^{\pm 1}$ helix (RNA binding) | 1R6R | RNA binding region packages viral genome |
| 74 | C | 44 | $\hat{I}^{\pm 1}$ helix (RNA binding) | 1R6R | RNA binding region packages viral genome |
| 91 | C | 61 | $\hat{I}^{\pm 2}$-$\hat{I}^{\pm 3}$ helices (dimerization) | 1R6R | |
| 150 | prM | 6 | pr domain<br>(chaperone<br>function) | 3J2P |  |
| 235 | prM | 91 | M ectodomain<br>(furin cleavage<br>site) | 3J2P |  |
| 337 | E | 27 | Domain I -<br>central | 1UZG, 3J2P, 3IRC |  |
| 342 | E | 32 | Domain I -<br>central | 1UZG, 3J2P, 3IRC |  |
| 433 | E | 123 | Domain I -<br>extension | 1UZG, 3J2P, 3IRC |  |
| 447 | E | 137 | Domain II -<br>dimerization | 1UZG, 3J2P, 3IRC |  |
| 448 | E | 138 | Domain II -<br>dimerization | 1UZG, 3J2P, 3IRC |  |
| 930 | NS1 | 127 | Wing domain | 4O6B |  |
| 931 | NS1 | 128 | Wing domain | 4O6B |  |
| 1375 | NS2B | 2 | N-terminal<br>transmembrane | 2FOM |  |
| 1376 | NS2B | 3 | N-terminal<br>transmembrane | 2FOM |  |
| 1476 | NS2B | 103 | C-terminal<br>transmembrane | 2FOM |  |
| 1480 | NS2B | 107 | C-terminal<br>transmembrane | 2FOM |  |
| 1664 | NS3 | 161 | Protease<br>domain | 2VBC,<br>2DV5 | Protease domain cleaves the viral<br>polyprotein |
| 1667 | NS3 | 164 | Protease<br>domain | 2VBC,<br>2DV5 | Protease domain cleaves the viral<br>polyprotein |
| 1675 | NS3 | 172 | Linker region | 2VBC, 2DV5 |  |

## References

1. Adikari, T. N., Riaz, N., Sigera, C., Leung, P., Valencia, B. M., Barton, K., . . . Rodrigo, C. (2020). Single molecule, near full-length genome sequencing of dengue virus. Scientific Reports, 10(1). doi:10.1038/s41598-020-75374-1

2. Adikari, T. N., Riaz, N., Sigera, C., Leung, P., Valencia, B. M., Barton, K., . . . Rodrigo, C. (2020). Single molecule, near full-length genome sequencing of dengue virus. Sci Rep, 10(1), 18196. doi:10.1038/s41598-020-75374-1

3. Avirutnan, P., Punyadee, N., Noisakran, S., Komoltri, C., Thiemmeca, S., Auethavornanan, K., . . . Malasit, P. (2006). Vascular leakage in severe dengue virus infections: a potential role for the nonstructural viral protein NS1 and complement. J Infect Dis, 193(8), 1078–1088. doi:10.1086/500949

4. Bhatt, S., Gething, P. W., Brady, O. J., Messina, J. P., Farlow, A. W., Moyes, C. L., . . . Hay, S. I. (2013). The global distribution and burden of dengue. Nature, 496(7446), 504–507. doi:10.1038/nature12060

5. Bull, R. A., Eltahla, A. A., Rodrigo, C., Koekkoek, S. M., Walker, M., Pirozyan, M. R., . . . Luciani, F. (2016). A method for near full-length amplification and sequencing for six hepatitis C virus genotypes. BMC Genomics, 17, 247. doi:10.1186/s12864-016-2575-8

6. Bull, R. A., Luciani, F., McElroy, K., Gaudieri, S., Pham, S. T., Chopra, A., . . . Lloyd, A. R. (2011). Sequential Bottlenecks Drive Viral Evolution in Early Acute Hepatitis C Virus Infection. PLOS Pathogens, 7(9), e1002243. doi:10.1371/journal.ppat.1002243

7. de Matos Simoes, R., & Emmert-Streib, F. (2011). Influence of Statistical Estimators of Mutual Information and Data Heterogeneity on the Inference of Gene Regulatory Networks. PLOS ONE, 6(12), e29279. doi:10.1371/journal.pone.0029279

8. Delport, W., Poon, A. F. Y., Frost, S. D. W., & Kosakovsky Pond, S. L. (2010). Datamonkey 2010: a suite of phylogenetic analysis tools for evolutionary biology. Bioinformatics, 26(19), 2455–2457. doi:10.1093/bioinformatics/btq429

9. Domingo, E., & Perales, C. (2019). Viral quasispecies. PLoS Genet, 15(10), e1008271. doi:10.1371/journal.pgen.1008271

10. Forns, X., Purcell, R. H., & Bukh, J. (1999). Quasispecies in viral persistence and pathogenesis of hepatitis C virus. Trends in Microbiology, 7(10), 402–410. 10.1016/S0966-842X(99)01590-5

11. Guo, C., Zhou, Z., Wen, Z., Liu, Y., Zeng, C., Xiao, D., . . . Yang, G. (2017). Global Epidemiology of Dengue Outbreaks in 1990-2015: A Systematic Review and Meta-Analysis. Front Cell Infect Microbiol, 7, 317. doi:10.3389/fcimb.2017.00317

12. Hu, B., Guo, H., Zhou, P., & Shi, Z.-L. (2021). Characteristics of SARS-CoV-2 and COVID-19. Nature Reviews Microbiology, 19(3), 141–154. doi:10.1038/s41579-020-00459-7

13. Huy, N. T., Van Giang, T., Thuy, D. H., Kikuchi, M., Hien, T. T., Zamora, J., & Hirayama, K. (2013). Factors associated with dengue shock syndrome: a systematic review and meta-analysis. PLoS Negl Trop Dis, 7(9), e2412. doi:10.1371/journal.pntd.0002412

14. Kosakovsky Pond, S. L., & Frost, S. D. W. (2005). Not So Different After All: A Comparison of Methods for Detecting Amino Acid Sites Under Selection. Molecular Biology and Evolution, 22(5), 1208–1222. doi:10.1093/molbev/msi105

15. Kosakovsky Pond, S. L., Poon, A. F. Y., Velazquez, R., Weaver, S., Hepler, N. L., Murrell, B., . . . Muse, S. V. (2019). HyPhy 2.5—A Customizable Platform for Evolutionary Hypothesis Testing Using Phylogenies. Molecular Biology and Evolution, 37(1), 295–299. doi:10.1093/molbev/msz197

16. Lindenbach, B. D., & Rice, C. M. (1997). trans-Complementation of yellow fever virus NS1 reveals a role in early RNA replication. J Virol, 71(12), 9608–9617. doi:10.1128/jvi.71.12.9608-9617.1997

17. Maduranga, S., Valencia, B. M., Sigera, C., Adikari, T., Weeratunga, P., Fernando, D., . . . Rodrigo, C. (2023). Genomic Surveillance of Recent Dengue Outbreaks in Colombo, Sri Lanka. Viruses, 15(7). doi:10.3390/v15071408

18. Miller, G. (1955). Note on the bias of information estimates. Information theory in psychology: Problems and methods.

19. Muñoz-Jordan, J. L., Sánchez-Burgos, G. G., Laurent-Rolle, M., & García-Sastre, A. (2003). Inhibition of interferon signaling by dengue virus. Proc Natl Acad Sci U S A, 100(24), 14333–14338. doi:10.1073/pnas.2335168100

20. Murrell, B., Moola, S., Mabona, A., Weighill, T., Sheward, D., Kosakovsky Pond, S. L., & Scheffler, K. (2013). FUBAR: A Fast, Unconstrained Bayesian AppRoximation for Inferring Selection. Molecular Biology and Evolution, 30(5), 1196–1205. doi:10.1093/molbev/mst030

21. Murrell, B., Wertheim, J. O., Moola, S., Weighill, T., Scheffler, K., & Kosakovsky Pond, S. L. (2012). Detecting Individual Sites Subject to Episodic Diversifying Selection. PLoS Genetics, 8(7), e1002764. doi:10.1371/journal.pgen.1002764

22. Pond, S. L. K., & Frost, S. D. W. (2005). Datamonkey: rapid detection of selective pressure on individual sites of codon alignments. Bioinformatics, 21(10), 2531–2533. doi:10.1093/bioinformatics/bti320

23. Pond, S. L. K., Frost, S. D. W., & Muse, S. V. (2004). HyPhy: hypothesis testing using phylogenies. Bioinformatics, 21(5), 676–679. doi:10.1093/bioinformatics/bti079

24. Rajapakse, S., Rodrigo, C., & Rajapakse, A. (2012). Treatment of dengue fever. Infect Drug Resist, 5, 103–112. doi:10.2147/idr.S22613

25. Real-time tracking of dengue virus evolution. (2025, 05-05-2025). Retrieved from https://nextstrain.org/dengue/all/genome

26. Riaz, N., Leung, P., Barton, K., Smith, M. A., Carswell, S., Bull, R., . . . Rodrigo, C. (2021). Adaptation of Oxford Nanopore technology for hepatitis C whole genome sequencing and identification of within-host viral variants. BMC Genomics, 22(1), 148. doi:10.1186/s12864-021-07460-1

27. Riaz, N., Leung, P., Bull, R. A., Lloyd, A. R., & Rodrigo, C. (2022). Evolution of within-host variants of the hepatitis C virus. Infect Genet Evol, 105242. doi:10.1016/j.meegid.2022.105242

28. Rodrigo, C., & Luciani, F. (2019). Dynamic interactions between RNA viruses and human hosts unravelled by a decade of next generation sequencing. Biochim Biophys Acta Gen Subj, 1863(2), 511–519. doi:10.1016/j.bbagen.2018.12.003

29. Sangkaew, S., Ming, D., Boonyasiri, A., Honeyford, K., Kalayanarooj, S., Yacoub, S., . . . Holmes, A. (2021). Risk predictors of progression to severe disease during the febrile phase of dengue: a systematic review and meta-analysis. Lancet Infect Dis, 21(7), 1014–1026. doi:10.1016/s1473-3099(20)30601-0

30. Santiago, G. A., Vergne, E., Quiles, Y., Cosme, J., Vazquez, J., Medina, J. F., . . . Muñoz-Jordán, J. L. (2013). Analytical and Clinical Performance of the CDC Real Time RT-PCR Assay for Detection and Typing of Dengue Virus. PLOS Neglected Tropical Diseases, 7(7), e2311. doi:10.1371/journal.pntd.0002311

31. Shannon, C. E. (1948). A Mathematical Theory of Communication. Bell System Technical Journal, 27(3), 379–423. 10.1002/j.1538-7305.1948.tb01338.x

32. Sigera, C., Rodrigo, C., De Silva, N. L., Weeratunga, P., Fernando, D., & Rajapakse, S. (2021). Direct costs of managing in-ward dengue patients in Sri Lanka: A prospective study. PLOS ONE, 16(10), e0258388. doi:10.1371/journal.pone.0258388

33. Sigera, P. C., Amarasekara, R., Rodrigo, C., Rajapakse, S., Weeratunga, P., De Silva, N. L., . . . Fernando, S. D. (2019). Risk prediction for severe disease and better diagnostic accuracy in early dengue infection; the Colombo dengue study. BMC Infectious Diseases, 19(1). doi:10.1186/s12879-019-4304-9

34. Weaver, S., Shank, S. D., Spielman, S. J., Li, M., Muse, S. V., & Kosakovsky Pond, S. L. (2018). Datamonkey 2.0: A Modern Web Application for Characterizing Selective and Other Evolutionary Processes. Molecular Biology and Evolution, 35(3), 773–777. doi:10.1093/molbev/msx335

35. World Health Organization. (2009). Dengue guidelines for diagnosis, treatment, prevention and control. New Delhi: WHO.

36. Xie, X., Gayen, S., Kang, C., Yuan, Z., & Shi, P. Y. (2013). Membrane topology and function of dengue virus NS2A protein. J Virol, 87(8), 4609–4622. doi:10.1128/jvi.02424-12

37. Yung, C. F., Lee, K. S., Thein, T. L., Tan, L. K., Gan, V. C., Wong, J. G. X., . . . Leo, Y. S. (2015). Dengue serotype-specific differences in clinical manifestation, laboratory parameters and risk of severe disease in adults, singapore. Am J Trop Med Hyg, 92(5), 999–1005. doi:10.4269/ajtmh.14-0628

38. Zhang, Y., Yin, Q., Ni, M., Liu, T., Wang, C., Song, C., . . . Ma, L. (2021). Dynamics of HIV-1 quasispecies diversity of participants on long-term antiretroviral therapy based on intrahost single-nucleotide variations. International Journal of Infectious Diseases, 104, 306–314. 10.1016/j.ijid.2021.01.015

